# Enhancing single-cell ATAC sequencing with formaldehyde fixation, cryopreservation, and multiplexing for flexible analysis

**DOI:** 10.1101/2024.11.20.624480

**Authors:** Tobias Hohl, Ulrike Boenisch, Thomas Manke, Laura Arrigoni

## Abstract

The assay for transposase-accessible chromatin using sequencing (ATAC-seq) revolutionized the field of epigenetics since its emergence by providing a means to uncover chromatin dynamics and other factors affecting gene expression. The development of single-cell (sc) applications in recent years led to an even deeper understanding of cell type specific gene regulatory mechanisms. One of the major challenges while running ATAC-seq experiments, bulk or sc, is the need for freshly collected cells for successful experiments. While various freezing methods have already been tested and established for bulk and sc ATAC-seq, quality metrics for preserved cells are rather poor or dependent on sampling time when compared to fresh samples. This makes it difficult to conduct all sorts of complex experiments i.e. with multiple conditions, patients, or time course studies. Especially, accounting for batch effects can be difficult if samples need to be processed at different time points of collection. We tackled this issue by adding a fixation step prior to the freezing method. The additional fixation step improved library quality and yield data comparable to fresh samples. The workflow was also tested on multiplexed sc ATAC experiments, set-up for cost-efficient low input sample handling. Sample cross-in, typically encountered in Tn5-based multiplex approaches, were tackled with a computational procedure specifically developed for this approach.

## Introduction

The assay for transposase-accessible chromatin using sequencing (ATAC-seq) has become a pivotal tool for studying chromatin accessibility, offering insights into gene regulation across various applications (Buenrostro et al., 2013). Despite its widespread use, a significant challenge remains: the requirement for fresh samples, which can introduce logistical hurdles and technical artifacts during sampling and preservation, affecting reproducibility and data quality (Grandi et al., 2022). Studies have shown that delays in sample processing and cryopreservation can distort chromatin accessibility profiles, leading to biased results (Massoni-Badosa et al., 2020). Additionally, the use of DMSO in cryopreservation, although largely used in ATAC-seq (Chen et al., 2018; Wong et al., 2023), can also introduce unpredictable effects on chromatin structure and gene expression.

To address these challenges, we explore a two-step procedure involving formaldehyde fixation combined with cryopreservation and flash freezing. This approach seeks to initially stabilize samples through formaldehyde treatment, effectively preventing further biological changes during collection and storage, while extending the applicability of ATAC-seq to archived samples. Previous studies have demonstrated the utility of formaldehyde in preserving chromatin structure for ATAC-seq (Chen et al., 2016), even in formalin-fixed paraffin-embedded (FFPE) samples (Zhang et al., 2022). Additionally, methodologies like FixNCut have shown the potential of using fixatives in preserving RNA integrity and reducing artifacts in single-cell assays (Jiménez-Gracia et al., 2024).

Our study builds on these findings by systematically evaluating the effects of formaldehyde fixation on single-cell ATAC-seq (scATAC-seq) using the 10x Genomics Chromium platform. We benchmark various preservation techniques, including formaldehyde fixation at several concentrations, followed by different freezing methods, to identify optimal conditions that maintain data quality comparable to fresh samples. We identified that the optimal condition involves fixing samples with a low concentration of formaldehyde (0.1%) followed by DMSO cryopreservation, yielding high quality data that best resembles the data obtained from fresh, untreated samples. This enhances robustness of library preparation protocols and provides practical guidelines for preserving samples in complex experimental designs, such as time-course studies with multiple conditions.

In addition to enhancing sample preservation, we have developed a streamlined transposase-based multiplexing strategy aimed at reducing costs and improving workflow efficiency. This method allows for the collection of samples at specific time points, facilitating high-throughput analysis using the 10x Genomics platform. By integrating sample-specific barcodes during transposition, our method simplifies both laboratory handling and data analysis, eliminating the need for additional sequencing libraries. Given the low formaldehyde concentration used, we envision that this method might be applicable to other Tn5-based studies, such as CUT&Tag and single-cell multiomic approaches.

In summary, our study presents a comprehensive approach to overcoming the limitations of fresh sample requirements in ATAC-seq, ensuring high-quality data generation while addressing cost concerns through multiplexing. This work paves the way for more flexible and robust study designs in chromatin accessibility research.

## Results

### Combining formaldehyde fixation with sample preservation methods yields good quality bulk ATAC-seq data

Using HepG2 as our model, we harvested cells and treated them with varying concentrations of formaldehyde (FA), ranging from 0.1% to 5%. This broad range was chosen to thoroughly investigate the effects of formaldehyde fixation on chromatin accessibility. While a 1% formaldehyde concentration is typically standard in chromatin assays such as ChIP-seq or Hi-C, higher concentrations have been reported to enhance signal strength in ChIP-seq experiments (Zaidi et al., 2017; Zhao et al., 2020). Therefore, we included higher concentrations to determine if similar improvements could be observed in ATAC-seq data quality. Following fixation, the cells were preserved using either cryopreservation or flash freezing to assess the impact of these preservation methods on data integrity.

ATAC-seq libraries were generated following the “Omni-ATAC” procedure (Corces et al., 2017) (see Methods) and sequenced and compared with publicly available data from ENCODE, where fresh untreated cells were used (Fig. 1a). Overall, the samples showed very comparable data quality. While all flash frozen samples had a FRIPs score of around 20%, this score was significantly higher for cryopreserved samples fixed with low FA concentrations, which scored ∼30% matching the ENCODE reference, but seemed to decrease with increasing FA concentration (Fig. 1b). Overall, higher FA concentration also yielded a higher fraction of mitochondrial read contents (Fig. 1c) and led to a noisier genome track signal (Suppl. Fig. 1), while the signal around ENCODE peak regions was very comparable between all conditions (Fig. 1e). We conclude that mild fixation (<5% FA) prior to preservation yields ATAC seq data of comparable quality as for reference data from fresh HepG2 cells.

**Fig. 1:**
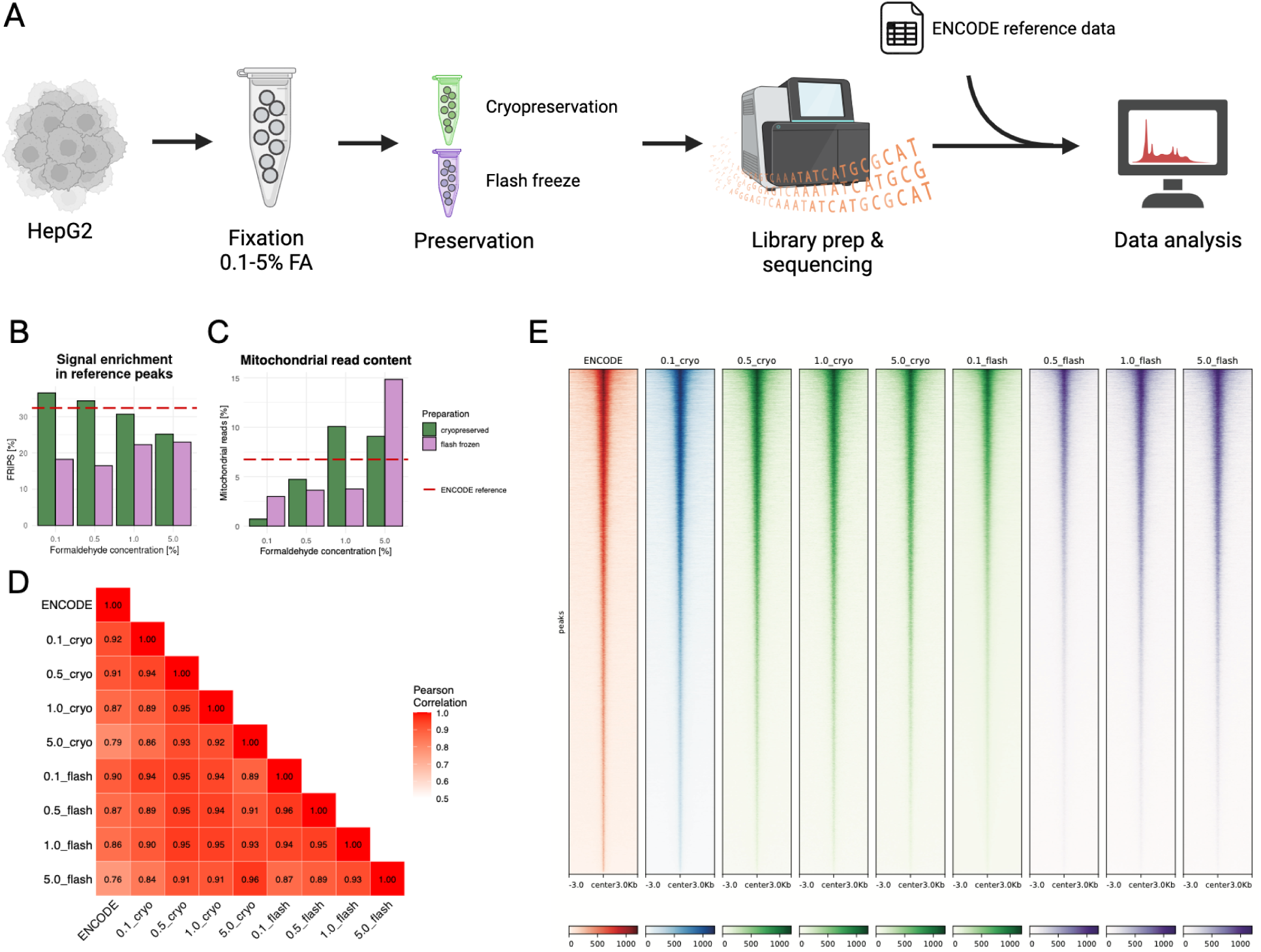
Coupling formaldehyde fixation with preservation methods yields good quality ATAC-seq data. **a** Experimental workflow. Created in BioRender. Arrigoni, L. (2024) https://BioRender.com/u59v773 **b-c** Quality metrics of various fixation and preservation combinations. **d** Pearson correlation heatmap between different conditions. **e** Heatmap showing signal enrichment around reference peaks for selected conditions.

### Moderate fixation combined with cryopreservation outperforms other treatments

Our analysis revealed that samples treated with higher concentrations of FA exhibited lower performance across several quality metrics. This trend is further supported by sample-to-sample correlation, where samples fixed with 5% FA demonstrated the weakest correlation with the reference data (Fig. 1d). On the other hand, the cryopreserved sample with 0.1% FA shows the best correlation with the reference (Fig. 1d). In addition, both fragment size distribution (Suppl. Fig. 2a) and PCA (Suppl. Fig. 2b) confirm the slightly better performance of cryopreservation with 0.1% FA. A transcription factor (TF) footprint analysis of HepG2-associated TF motifs (Huang et al., 2020) also shows that increasing FA concentration appears to reduce the strength of the footprint signal (Suppl. Fig. 3). From this, we conclude that 0.1% FA is the best performing FA concentration and therefore would be used as the concentration for the single-cell experiments stated below.

To avoid biases and common uncertainties in the peak calling, we have used only ENCODE reference peaks above. If we add peaks calls from our lightly fixed (0.1 % FA) and cryopreserved data, we also find a large overlap (∼70% of peaks, SFig 2C). Inspection of the heatmap (Sfig 2D) shows a very consistent signal also for those regions that were not jointly called as peaks - this reflects threshold effects and possible differences in statistical power. Importantly the fixation does not appear to introduce artificial peaks with respect to the reference data produced from fresh, untreated samples.

**Fig. 2:**
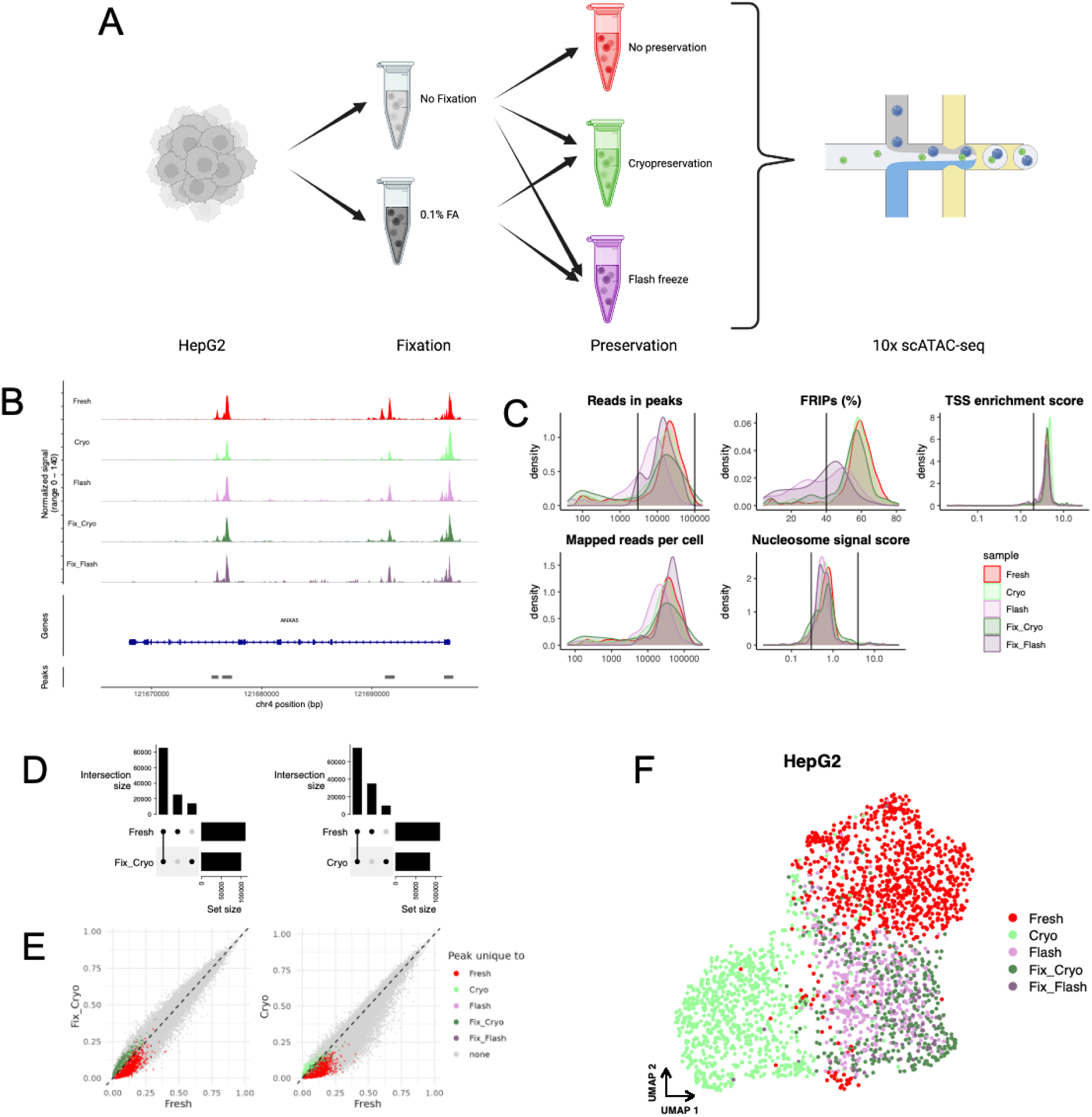
Microfluidics based scATAC-seq can be performed with fixed and preserved samples. **a** Experimental workflow. Created in BioRender. Arrigoni, L. (2024) https://BioRender.com/s08b492 **b** Genome tracks showing the signal per sample on a select region. **c** Single-cell data quality metrics. **d** Upset plots comparing peak sets called per sample. **e** Correlation plots of fractions of cells per sample that show signal in the respective peak region. **f** UMAP representation of HepG2 scATAC-seq data.

### Evaluating preservation methods for single-cell ATAC-seq applications

Building on our bulk analysis findings, we integrated the fixation step into the 10x Genomics single-cell workflow to assess its effectiveness. We conducted a comparative study of scATAC-seq on HepG2 cells under various conditions: fresh, cryopreserved, flash-frozen, 0.1% FA with cryopreservation, and 0.1% FA with flash freezing (Fig. 2a and Methods). We evaluated these five distinct conditions, accompanied by comprehensive quality control assessments.

While most quality metrics we examined were quite comparable across all conditions, we observed that flash-freezing tends to decrease the signal-to-noise ratio. This is reflected by a reduced FRIP score in our flash-frozen and 0.1% formaldehyde followed by flash-freezing samples, whereas cryopreserved samples more closely resemble the fresh untreated sample (Fig 2c). This is noticeable also as a higher background in a representative IGV plot, while the overall shape of pseudo-bulk ATAC tracks is very comparable (Fig 2b). Additionally, flash freezing seems to overall lower TSS accessibility whereas cryopreservation does not show this trend (Suppl. Fig. 4b). In examining the fragment size distributions, we observed that flash-frozen or cryopreserved samples lacking fixation lost the distinct nucleosomal pattern. However, the introduction of a fixation step appeared to restore this characteristic feature (Suppl. Fig. 4a).

At cellular level, differences between the protocols can be visualized as UMAP embedding (Fig 2e, Suppl. Fig. 4c). We study those differences by various methods. Firstly, we look at the loadings of the components of the Latent Semantic Indexing (LSI) based dimension reduction that provides the basis for the UMAP embedding (Suppl. Fig. 5a). After identifying the LSI component responsible for sample separation, we look at the loadings and further investigate the top correlated regions (Suppl. Fig. 5b-e). While these regions show an increased signal within the Fresh sample, all samples show signal in the respective regions. Only the fixed sample followed by flash-freezing features a noisier and more sparse track (Suppl. Fig. 5e).

Subsequently, we look at differentially accessible regions (DARs) per protocol and discover that the amount of DARs is highest for the untreated, fresh sample (Suppl. Fig. 6a). While DAR sizes are very comparable across all groups (Suppl. Fig. 6b), we looked at the top DARs for each protocol and discovered differences in signal strength between samples, but no absence of signal within any group (Suppl. Fig. 6c).

Sample-specific peak calling yielded a comprehensive peak set that we used for a correlation analysis (Suppl. Fig. 7a, Fig. 2d). We compared the fraction of cells with signal in respective peak regions (Suppl. Fig. 7b, Fig. 2d) and discovered that preservation led to an overall decrease in cell fractions with reads in the majority of peaks, while the addition of fixation before preservation rescued this behavior. Furthermore, differentially called peaks were expectedly found in low fractions of cells, suggesting that the observed differences are mostly because of differences in statistical power for protocols with different cell numbers and sequencing depth, and ultimately threshold effects of the peak caller, rather than true biological differences.

While our data indicates the feasibility of different preservation techniques in the context of preparing scATAC-seq libraries using the 10x Chromium workflow, it also suggests that adding a fixation step with 0.1% FA before preservation increases data quality and rescues signals that might get lost otherwise. Overall, combining moderate fixation (0.1% FA) with cryopreservation yields data that resembles the untreated fresh sample best.

### Dealing with barcode hopping in tagmentation-based scATAC-seq sample multiplexing

Following earlier work (Lareau et al., 2019), we developed a multiplexing approach for scATAC-seq experiments that uses Tn5 enzymes pre-loaded with custom-made barcodes. In brief, the tagmentation is performed per sample, where the Tn5s insert the sample-specific barcodes into the cut sites. After tagmentation, the samples are pooled for library preparation and sequencing following standard protocols from 10x genomics and Illumina (see Methods). The data can be demultiplexed using the sample specific barcodes (Fig. 3a). We applied this workflow and pooled ten individually labeled samples for library preparation (Fig. 3b). Notably, only around 20% of cell barcodes were unique to a certain sample, while the rest appeared in two or more samples (Fig. 3c). It is important to note that we adhered to 10x Genomics recommendations on nuclei amount to load, so the multiplet rate is not derived by overloading. Given an anticipated doublet ratio of approximately 8%, we expected a similar percentage of barcodes to be present in multiple samples, but the observed number was unexpectedly high. Another study utilizing a very similar multiplexing approach does also report this effect through experiments with cell mixtures from separate species and an increase in inter-species doublet formation (Zhang et al., 2024). They state this issue derives from free floating, unbound Tn5 inserts that can enter GEMs after pooling and serve as primers for library preparation, thus introducing erroneous sample barcodes.

**Fig. 3:**
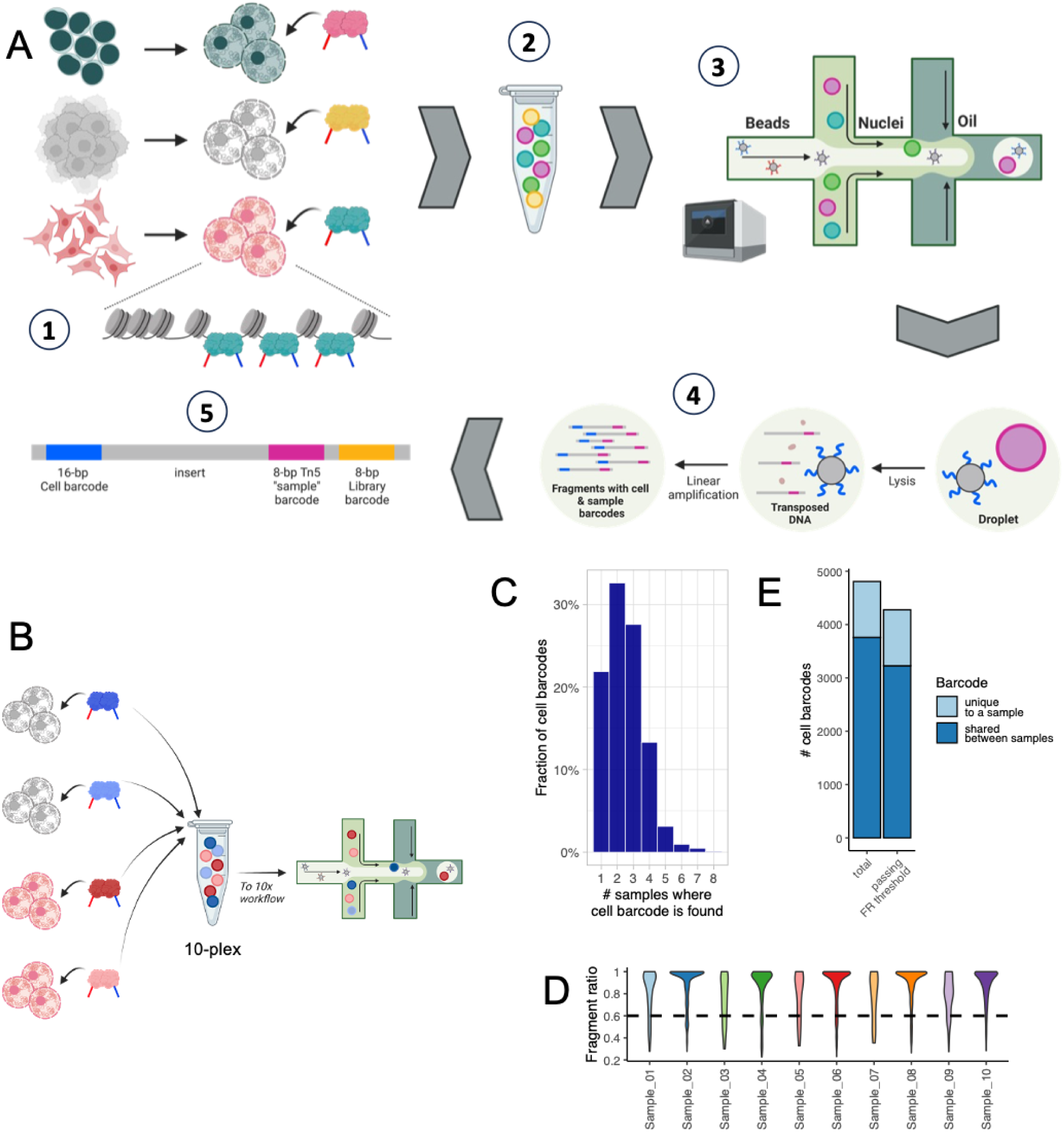
Multiplexing approach for scATAC-seq. **a** Tn5 barcoding workflow. 1 – Isolated nuclei are transposed using hyperactive Tn5 with pre-loaded sample barcode constructs. 2 – After Tn5 inactivation, samples are pooled. 3 & 4 – 10x Genomics barcoded microbeads and transposed nuclei are encapsulated within droplets, also called GEMs (Gel beads in EMulsion). Cells are lysed inside GEMs and transposed DNA fragments are linearly amplified. At this stage single cell barcodes are incorporated in the transposed fragments. 5 – PCR amplification of final fragments. The fragments include cell and library barcodes as well as the sample barcode that was introduced via Tn5 tagmentation. **b** Experimental workflow for testing the multiplexing protocol. **c** Cell barcode appearance across multiplexed samples. **d** Fragment ratio distributions across samples. Cell barcodes were assigned to the sample with the most fragments. **e** Cell barcode retainment after FR thresholding. Panels a&b: Created in BioRender. Arrigoni, L. (2024) https://BioRender.com/j56f499

The presence of a cellular barcode in multiple samples is a concern in terms of proper data demultiplexing, but rather than using a simple presence-absence pattern we make cell-sample assignments based on the normalized fragment counts N seen for cellular barcode c in sample s. Briefly, we will assign a cellular barcode c to sample s if more than 60% of all fragments with barcode c derive from sample s. This metric, which we call Fragment Ratio r can be calculated as follows:

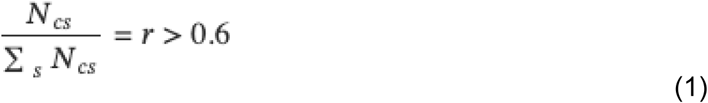

Fragment ratios for all sample and cell barcode combinations are shown in Suppl. Fig. 8a-b. For most cell barcodes, the highest contributing sample accounts for the majority of fragments, resulting in them passing the applied threshold of 60% (Fig. 3d). Through this threshold, we are able to assign more than 88% of all cellular barcodes to samples, compared to the 22% from a simple incidence (Fig. 3e).

Despite the challenge of severe barcode hopping, our bioinformatics approach effectively mitigates this issue, ensuring minimal data loss and enhancing the accuracy of sample assignment.

## Discussion

Disentangling the timing of sample collection from preparation is crucial for optimizing various experimental setups. In our study, we demonstrate that integrating formaldehyde fixation with cryopreservation in microfluidics-based single-cell ATAC-seq experiments produces results comparable to those from fresh samples. This method preserves a strong signal-to-noise ratio and maintains overall data integrity.

We offer a comprehensive benchmarking of sample preservation techniques for scATAC-seq library preparation, comparing two freezing methods: flash freezing and cryopreservation, both with and without prior moderate fixation (0.1% FA). Although standard batch correction algorithms can mitigate differences between protocols, it is advisable to maintain consistency by using the same protocol throughout a project. In large-scale projects, logistical constraints may prevent the timely processing of fresh samples. Our findings indicate that protocols involving fixation followed by preservation are suitable for standardized treatment and high reproducibility. Notably, fixed cryo-preserved samples exhibit the closest resemblance to fresh samples in terms of signal-to-noise ratio and overall similarity.

We also anticipate that our method could be applicable to scRNA-seq and single-cell multiome protocols. Incorporating light formaldehyde fixation before cryopreservation may help eliminate DMSO-induced artifacts on gene expression (Verheijen et al., 2019). Additionally, using light rather than strong fixation could obviate the need for probe methods, as suggested by recent studies (Llora-Batlle et al., 2024), thereby broadening the method’s applicability across different organisms.

Additionally, we have developed a sample multiplexing strategy that might leverage the potential of higher capacity platforms such as Chromium X, which can currently process up to 20,000 cells for scRNA-seq and may extend to scATAC-seq in the future. By collecting and fixing samples on different days, particularly those with low cell counts (e.g., fewer than 5,000 cells), researchers can fully utilize the capacity of droplet based methods. This approach maximizes the cost-effectiveness of high-throughput analysis tools, enabling more efficient and economical research workflows.

The multiplexing of scATAC-seq experiments has recently garnered significant attention, with various protocols emerging (Bera et al., 2023; Han et al., 2022; Wang et al., 2021). Although our approach was independently developed to reduce sequencing costs per reaction for small-scale experiments, we encountered cross-labeling of samples. A recent study presenting a similar multiplexing approach reports analogous issues (Zhang et al., 2024). When multiplexing samples from different species, inter-species doublet rates increased significantly. The authors suggest that this likely results from free-floating Tn5 adapters after sample pooling, which can interact with DNA fragments from cells not belonging to the respective sample barcode, thus affecting library preparation. While they propose a modified protocol involving the introduction of a reverse primer for exponential amplification during library preparation, our *in silico* approach filters using the fragment ratio without needing protocol modifications and works with moderate data loss (12%).

In conclusion, we provide a multiplexed scATAC-seq workflow that allows for flexible sample collection, enabling time-course experiments to be processed within a single 10x scATAC-seq reaction. This method eliminates batch effects arising from different sampling and library preparation experiments due to streamlined preservation and parallel processing.

## Methods

### Cell culture

Human HepG2 cells (ATCC, HB-8065) were cultured in EMEM (Sigma-Aldrich M5650) supplemented with 10% Fetal Bovine Serum (Sigma-Aldrich, F7524), 2 mM GlutaMAX (Gibco, 35050038), and 1mM sodium pyruvate (Gibco, 11360070). Mouse EL4 cells (ATCC, TIB-39) were grown in high glucose DMEM (Gibco, 10569010) supplemented with 10% horse serum (Gibco, 26050070). Mouse 3T3-L1 cells (ATCC) were grown in high glucose DMEM (Gibco, 10569010) supplemented with 10% Fetal Bovine Serum (Sigma-Aldrich, F7524). All cells were cultured in an incubator at 37°C with 5% CO_2_. Cell passages were performed by trypsinization.

### Preservation

#### Formaldehyde fixation

Fixative buffer was prepared by diluting a freshly opened vial of 16% Formaldehyde (CST, 12606) to a concentration of 0.1% to 5% in serum-free DMEM (Gibco, 10569010). Up to 3 million cells in suspension were transferred into 1.5 ml tubes and pelleted at 300 × g for 5 minutes. After removing the media, pellets were resuspended in 1 ml of fixative buffer and incubated at room temperature for 5 minutes, with tubes inverted every 1-2 minutes to prevent cell sedimentation. The glycine step was omitted, and cells were directly pelleted at 300 × g for 5 minutes. The supernatant was discarded appropriately. Fixed cells were washed in 1 ml of DMEM supplemented with 0.1% BSA, pelleted again, and either snap-frozen by submerging the tubes in liquid nitrogen or cryopreserved. Both snap-frozen and cryopreserved cells were stored at -70°C for long-term preservation.

#### Cryopreservation and thawing

Fixed or fresh cell pellets containing up to 3 million cells were resuspended in 1 ml of room temperature cryopreservation buffer (10% DMSO, 90% Fetal Bovine Serum). The vials were quickly transferred into a Mr. Frosty container equilibrated at room temperature and then placed in a -70°C freezer for at least overnight storage. For thawing, cryopreserved cells were quickly defrosted for up to 2 minutes in a thermoblock at 37°C, removing the vials when only a small ice crystal remained. Cell suspensions in DMSO were mixed and added dropwise to a 5 ml tube containing 4 ml of 1% BSA in PBS. Tubes were mixed by inversion and centrifuged at 300 × g for 5 minutes. Cells were then resuspended in the desired volume of fresh PBS/BSA and placed on ice.

### Nuclei isolation

Cells in suspension were pelleted at 300 × g for 5 minutes, resuspended in fresh media, and placed on ice. Using a hemocytometer, cells were counted, and up to 2 million cells were transferred into 1.5 ml tubes and pelleted at 300 × g for 5 minutes at 4°C. Cells and nuclei were permeabilized in 100 μl of lysis buffer (10 mM Tris-HCl, pH 7.4, 10 mM NaCl, 3 mM MgCl2, 0.1% Igepal CA-630, 0.1% Tween-20, 0.01% Digitonin, 1% BSA) and incubated for 5 minutes on ice. One milliliter of ice-cold wash buffer (10 mM Tris-HCl, pH 7.4, 10 mM NaCl, 3 mM MgCl2, 0.1% Tween-20, 1% BSA) was added to the cell suspension and mixed by inversion.

For bulk ATAC experiments, an aliquot containing 50,000 nuclei was separated into a 1.5 ml tube, centrifuged at 500 × g for 5 minutes at 4°C, and resuspended in the transposition mix as detailed in the protocols below.

For single-cell ATAC protocols, nuclei in wash buffer were centrifuged at 500 × g for 5 minutes at 4°C. Supernatants were immediately discarded, and pellets were resuspended in 1x Nuclei Dilution Buffer (10x Genomics, 2000207) to achieve a nuclei concentration of 3,200 nuclei/μl.

### Barcode design and transposome assembly

#### Tn5 barcode design

We developed a method to determine suitable sample barcodes that could be used for scATAC-seq multiplexing (see github repository). Given the parameters barcode length, minimum and maximum GC content, minimum distance between barcodes, the method to determine sequence distances (Hamming or Levenshtein), and an optional predefined barcode, all possible barcodes are generated and subsequently filtered for GC content, no base triplication, only single base duplication, and representation of all bases. If no barcode is predefined, a barcode with a distance to its reverse complement passing the threshold is randomly selected from the filtered list. Otherwise the predefined barcode is used as the starting point. The remaining barcode list is filtered for elements and their reverse complement passing the distance threshold to the starting barcode. Then this step is repeated - a new barcode is sampled from the remaining list, and the list is filtered using distances of barcodes and their reverse complements to the sampled barcode - until no barcodes are remaining. The result is a list of suitable sample barcodes with a defined minimum distance between each other.

All Tn5 8-nucleotide barcodes used for this study were designed to maximize sequence diversity while ensuring a 50% GC content.

#### Transposase loading

Oligonucleotides were purchased from IDT, with HPLC purification selected for Mosaic End adapters A and B (ME-A and ME-B), and standard desalting for the Mosaic End reverse (ME-rev). A comprehensive list of all barcodes located in the ME-B oligos is provided in Additional File 1. Lyophilized ME-A, ME-B and ME-rev oligos were resuspended in Annealing Buffer (40mM Tris-HCl pH 8.0, 50mM NaCl) to a stock concentration of 100 μM. ME-A and ME-B oligos were annealed to the complementary common ME-rev oligo. For annealing, ME-A (or ME-B) oligo was mixed with ME-rev at 1:1 ratio and incubated in a thermocycler at 95 °C for 5 minutes. The mixture was then cooled to 65°C at a rate of -1°C/second, incubated at 65°C for 5 minutes, and further cooled to 4°C at the same rate.

To assemble the transposome, 5 μl of each annealed A and B transposome were combined with 10 μl of unloaded Tagmentase (Diagenode, C01070010), briefly vortexed, and incubated at 23°C for 30 minutes in a thermocycler. Subsequently, 10 μl of glycerol were added to the assembled transposase for storage at -20°C. The optimal dilution ratio of the stock Tn5 should be determined experimentally.

### Custom transposase titration using bulk ATAC-seq

To determine the optimal concentration of custom Tn5, libraries were prepared following the Omni ATAC-seq protocol with minor modifications and compared to those obtained using Illumina or 10x Genomics Tn5.

Nuclei suspension aliquots containing 50,000 nuclei were centrifuged at 500 × g for 5 minutes at 4°C. Each pellet was then resuspended in 47.5 μl of transposition mix (comprising 25 μl of Diagenode 2x Tn5 reaction buffer, 0.5 μl of 1% Digitonin, 0.5 μl of 10% Tween-20, 16.5 μl of PBS, and 5 μl of water) and placed on ice. Two-fold serial dilutions of the annealed Tn5 were prepared by diluting the stock Tn5 in Transposase dilution buffer (Diagenode, C01070011). To initiate transposition, 2.5 μl of custom diluted Tn5 (1:2 and 1:4), undiluted stock, ATAC enzyme B (10x Genomics, 2000265/72), or Illumina TDE1 Tagment enzyme (20034197) were added to each reaction, mixed by pipetting, and incubated for 30 minutes at 37°C in a thermocycler.

DNA was purified using the MinElute PCR purification kit (Qiagen, 28004) according to the manufacturer’s instructions, with a final elution volume of 21 μl. For PCR amplification, 10 μl of each transposed DNA was mixed with 2.5 μl of 25 μM i5 primer (same primers as dual index Cut&Tag libraries (Buenrostro et al., 2015), see also Additional File 1), 2.5 μl of 25 μM i7 primer (for 10x Genomics/Illumina Tn5, the Buenrostro 2015 list was used (Buenrostro et al., 2015); for custom Tn5, see Additional File 2), and 25 μl of 2x NEBNext Ultra II Q5 Master Mix (NEB, M0544). The mixture was amplified (72°C for 5 min, 98°C for 30 s, followed by 10 cycles of 98°C for 10 s, 63°C for 30 s, 72°C for 1 min, hold at 4°C).

Final libraries were cleaned using Ampure XP at a 0.8x ratio (40 μl of beads per 50 μl of reaction) and eluted in 25 μl of EB (Qiagen). Libraries were quantified using the Qubit High Sensitivity DNA Assay (Invitrogen, Q32851) and inspected for size distribution using a Fragment Analyzer with the NGS 1-6000 bp hs DNA kit. The optimal concentration of custom Tn5 for the single-cell assay was determined by ensuring the closest similarity in library size distribution to those obtained using Illumina or 10x Genomics Tn5.

### Multiplexed bulk ATAC sequencing

A pellet containing 50,000 nuclei was resuspended in 50 μl of transposition mix, which included 25 μl of Diagenode 2x Tn5 reaction buffer, 0.5 μl of 1% Digitonin, 0.5 μl of 10% Tween-20, 16.5 μl of PBS, 5 μl of water, and 2.5 μl of Custom Tn5 at a 1:4 dilution. Barcoded Tn5 were selected to ensure no barcode overlap after pooling. The mixture was thoroughly mixed by pipetting and incubated for 30 minutes at 37°C in a thermocycler, with the lid set at 50°C.

Transposition was quenched by adding 50 μl of 2x Tn5 stop solution (40 mM EDTA, 2 mM spermidine) to each reaction, followed by incubation at 37°C for 15 minutes in a thermocycler. The reactions were then cooled and pooled in a 1.5 ml tube on ice. The pooled cells were centrifuged at 500 × g for 5 minutes at 4°C and resuspended in 100 μl of buffer EB (Qiagen).

For fixed samples, prior to purification, samples were de-crosslinked at 68°C for 30 minutes in a thermocycler, with the lid set at 85°C.

DNA was purified using the MinElute PCR purification kit (Qiagen, 28004) according to the manufacturer’s instructions, with a final elution volume of 21 μl. For PCR amplification, 10 μl of each transposed DNA was mixed with 2.5 μl of 25 μM i5 primer (same primers as dual index Cut&Tag libraries (Buenrostro et al., 2015), see also Additional File 1), 2.5 μl of 25 μM i7 primer (MUXscATAC_i7_n primer, see Additional File 1), and 25 μl of 2x NEBNext Ultra II Q5 Master Mix (NEB, M0544). The mixture was amplified (72°C for 5 min, 98°C for 30 s, followed by 10 cycles of 98°C for 10 s, 63°C for 30 s, 72°C for 1 min, hold at 4°C).

Final libraries were cleaned using Ampure XP at a 0.8x ratio (40 μl of beads per 50 μl of reaction) and eluted in 25 μl of EB (Qiagen).

### 10x Genomics single-cell ATAC v2 workflow

Five microliters of permeabilized nuclei suspension, at a concentration of 3,200 nuclei/μl, were mixed with 7 μl of ATAC Buffer B (10x Genomics, 2000193) and 3 μl of ATAC Enzyme B (10x Genomics, 2000265/72). The transposition reaction was incubated in a thermocycler at 37°C for 30 minutes, with the lid set at 50°C. The single-cell ATAC v2 protocol was then continued from step 2.1 of the manufacturer’s manual (Chromium Next GEM Single Cell ATAC Reagent Kits v2 User Guide, CG000496 rev B). Final libraries were PCR-amplified using 7 cycles, as recommended for a target recovery of 10,000 cells.

### Multiplexed single-cell ATAC sequencing

A 5 μl aliquot of permeabilized nuclei suspension, at a concentration of 3,200 nuclei/μl, was added to 8.5 μl of transposition mix for multiplexed scATAC (comprising 7.5 μl of 2x Diagenode Tn5 reaction buffer and 1.5 μl of 10x Genomics 1x Nuclei Dilution Buffer) on ice. To each transposition reaction, 1 μl of custom barcoded Tn5 at a 1:4 dilution was added, ensuring the selection of Tn5 barcodes to avoid overlap after pooling. All reactions were mixed by pipetting and incubated in a thermocycler at 37°C for 30 minutes, with the lid set at 50°C.

Transposition was quenched by adding 1 μl of 300 mM EDTA to each reaction, followed by incubation at 37°C for 15 minutes in a thermocycler. The reactions were then cooled and pooled in a 1.5 ml tube on ice. The pooled nuclei were centrifuged at 500 × g for 5 minutes at 4°C and washed once with 500 μl of 1x Nuclei Dilution Buffer (10x Genomics, 2000207). Transposed cells were re-counted and diluted using 1x Nuclei Dilution Buffer to a concentration of 1,040 nuclei/μl.

Fifteen microliters of the diluted cell suspension were aliquoted into a PCR tube strip on ice for loading into the Chromium Controller, with a total of 15,600 cells loaded according to 10x Genomics recommendations. The single-cell ATAC v2 protocol was then continued from step 2.1 of the manufacturer’s manual (Chromium Next GEM Single Cell ATAC Reagent Kits v2 User Guide, CG000496 rev B).

For final PCR amplification at step 4.1 of the User Guide, the "individual single index set N" was replaced with 2.5 μl of 25 μM custom oligo (MUXscATAC_i7_n, sequences in Additional File 1), selecting library indices appropriately to prevent equal index assignment. Library amplification was performed using 9 PCR cycles.

### Library quality assessment and sequencing

Libraries were quantified using the Qubit High Sensitivity DNA Assay (Invitrogen, Q32851), and size distribution was visualized by capillary electrophoresis using the Fragment Analyzer with the NGS 1-6000 bp High Sensitivity DNA kit. DNA concentrations were adjusted based on the percentage of library fragments between 150 to 1200 bp, as determined by smear analysis. Library molarities were calculated using the adjusted DNA concentration and the average fragment size from the smear analysis.

Libraries were pooled, cleaned of adapter dimers, denatured according to Illumina guidelines, and sequenced paired-end with a read length of 16x50x50x32 bp (i5 index, R1, R2, i7 index; see Additional File 3 for 32-bp i7 index sequences) on a NovaSeq 6000 instrument. This setup was used for sequencing the single-cell and Tn5 sample barcodes along with the insert. In cases where only the Tn5 inserted barcode is desired to be read in i7, libraries can be sequenced with a read length of 16x50x50x32 bp. The 8-bp and 32-bp index list is provided in Additional File 1.

### BCL conversion and demultiplexing

BCL files were converted to fastq format using bcl2fastq2. Demultiplexing was performed using either Illumina p7 barcodes or sample barcodes introduced via multiplexing.

### Bulk ATAC-seq data analysis

#### Trimming & mapping

Demultiplexed fastq files were used to run the DNA-mapping pipeline from snakePipes (v2.7.2) (Bhardwaj et al., 2019) with default parameters (for reference, see github repo). Briefly, ATAC cut sites were trimmed from uniquely mapped pairs (mapq > 2) using cutadapt with parameters “--trimmerOptions -a nexteraF=CTGTCTCTTATA -A nexteraR=CTGTCTCTTATA” (added in DNA-mapping call). Mapping was performed on the human genome version GRCh38 using Bowtie2 and PCR duplicates were removed using samtools.

#### Further analyses

The output of the DNA-mapping pipeline was used for further analysis. For all peak-dependent analyses, the peak set of the reference ENCODE data was used. Most of the processing was conducted using the deepTools suite (v3.5.5) (Ramírez et al., 2016). Parameters for each step can be found on the github repository. Briefly, Coverage tracks were created using bamCoverage with RPKM normalization and a bin size of 1bp. Fragment size distribution data was generated using bamPEFragmentSize. FRIPs was determined using plotEnrichment. Both PCA (via plotPCA) and correlation (via plotCorrelation) analyses were performed using the output of MultiBigWigSummary. Peak coverage heatmaps were generated using computeMatrix and plotHeatmap. Genome tracks were plotted using pyGenomeTracks (Lopez-Delisle et al., 2021) with the generated bigwig files. The respective configuration files can be found on the github repository. Peaks for the 0.1% FA cryo sample were called using MACS2 with standard parameters. Bedtools2 was used for overlapping with the reference peak set, and the results were visualized using the UpSetR library. The TOBIAS package was used for TF motif footprinting analysis (Bentsen et al., 2020). Plots were generated using R (v4.3.3) and the tidyverse, ggbeeswarm, and ggstar packages. The respective scripts can be found in the repository.

### scATAC-seq data analysis

#### Preprocessing

Demultiplexed fastq files were processed by running the count pipeline of the CellRanger-ATAC software (v2.1.0) by 10x Genomics on each individual sample. Briefly, the pipeline filters all reads and aligns them to the reference genome. It demultiplexes cellular information using the cell barcode, identifies transposase cut sites, calls accessible peaks across the sample, and calls cells by determining high quality cell barcodes rich in signal-specific fragments.

The peak set used for downstream analysis was generated by running CellRanger-ATAC (v2.1.0) on publicly available HepG2 scATAC-seq data (ENCODE experiment ENCSR398OHC, R1 ENCFF074MZG, R2 ENCFF971PQX, cell barcode ENCFF084OJB).

#### Analysis

The fragment files and cell calling information provided by the results of the cellranger-ATAC count pipeline were used to input the single-cell information to the R-based Signac framework (v1.13) (Stuart et al., 2021) that served for most of the downstream analyses (see github repository). Briefly, count matrices were produced using the previously generated reference peakset and the sample-specific fragment files and cell calling information. After merging the count matrices, nucleosome signal and TSS enrichment scores are calculated. QC is performed by evaluating and filtering for FRIPS, reads in peaks, TSS enrichment and nucleosome signal scores. Subsequently, dimension reduction is performed by employing TF-IDF and SVD to generate LSI components. These serve as the basis for non-linear dimensionality reduction using UMAP. Differentially accessible regions were called using FindAllMarkers. Sample specific peaks were called using CallPeaks after accounting for sequencing depth differences that could significantly affect peak calling by subsampling to similar cell numbers. Resulting peak sets were compared using bedtoolsr and the UpSet function from the ComplexHeatmaps package. The resulting peak sets were combined and used to create a count matrix for all samples. This count matrix was binarized and then used to determine the fraction of cells per sample that show signal in each respective region / peak.

### Analysis of multiplexed scATAC-seq data

The data was preprocessed using the cellranger-ATAC count pipeline and read into the Signac framework, as previously described. To determine cross contamination of sample specific multiplexing barcodes, the occurrence of cell barcodes across samples was determined. We then developed a method to bioinformatically allocate unique cell barcodes to the respective samples. We make cell-sample assignments based on the normalized fragment counts seen for cellular barcode c in sample s. Briefly, we will assign a cellular barcode c to sample s if more than 60% of all fragments with cell barcode c derive from sample s (see Formula (1)). Downstream analysis followed the pipeline described above.

## Data availability

Datasets generated during this study are available in the Gene Express Omnibus repository under accession numbers GSE281808 for bulk ATAC-seq, and GSE281809 for multiplexed single-cell ATAC-seq of HepG2 cells. Bulk ATAC-Seq reference data was downloaded from ENCODE (Experiment ENCSR042AWH, library ENCLB324GIU).

## Code availability

Pipelines and scripts used to run analyses and generate figures in this study are available on github: https://github.com/TobiasHohl/scATAC-MUX-preservation.

## Acknowledgements

This work was funded and supported by the Deutsche Forschungsgemeinschaft (DFG, German Research Foundation) - 322977937/GRK2344. We would like to thank the deep sequencing core facility at the Max-Planck-Institute of Immunobiology and Epigenetics for their support in sequencing the libraries for this study. We also appreciate the assistance from the bioinformatics core facility, especially Ward Deboutte, for his help with data preprocessing, demultiplexing, and data submission.

## Authors and affiliations

Max-Planck-Institute of Immunobiology and Epigenetics, Freiburg Tobias Hohl, Ulrike Boenisch, Thomas Manke, Laura Arrigoni Faculty of Biology, Freiburg University, Freiburg Tobias Hohl

## Contributions

L.A. and T.H. designed the study. L.A. performed the experiments and produced the sequencing libraries. T.H. performed data processing and analysis, and developed the necessary algorithms. U.B. and T.M. supervised the work. T.H. and L.A. designed the figures. T.H. drafted the manuscript. All authors discussed the results and contributed to editing the manuscript.

## Corresponding author

Laura Arrigoni

---

**Suppl. Fig. 1:**
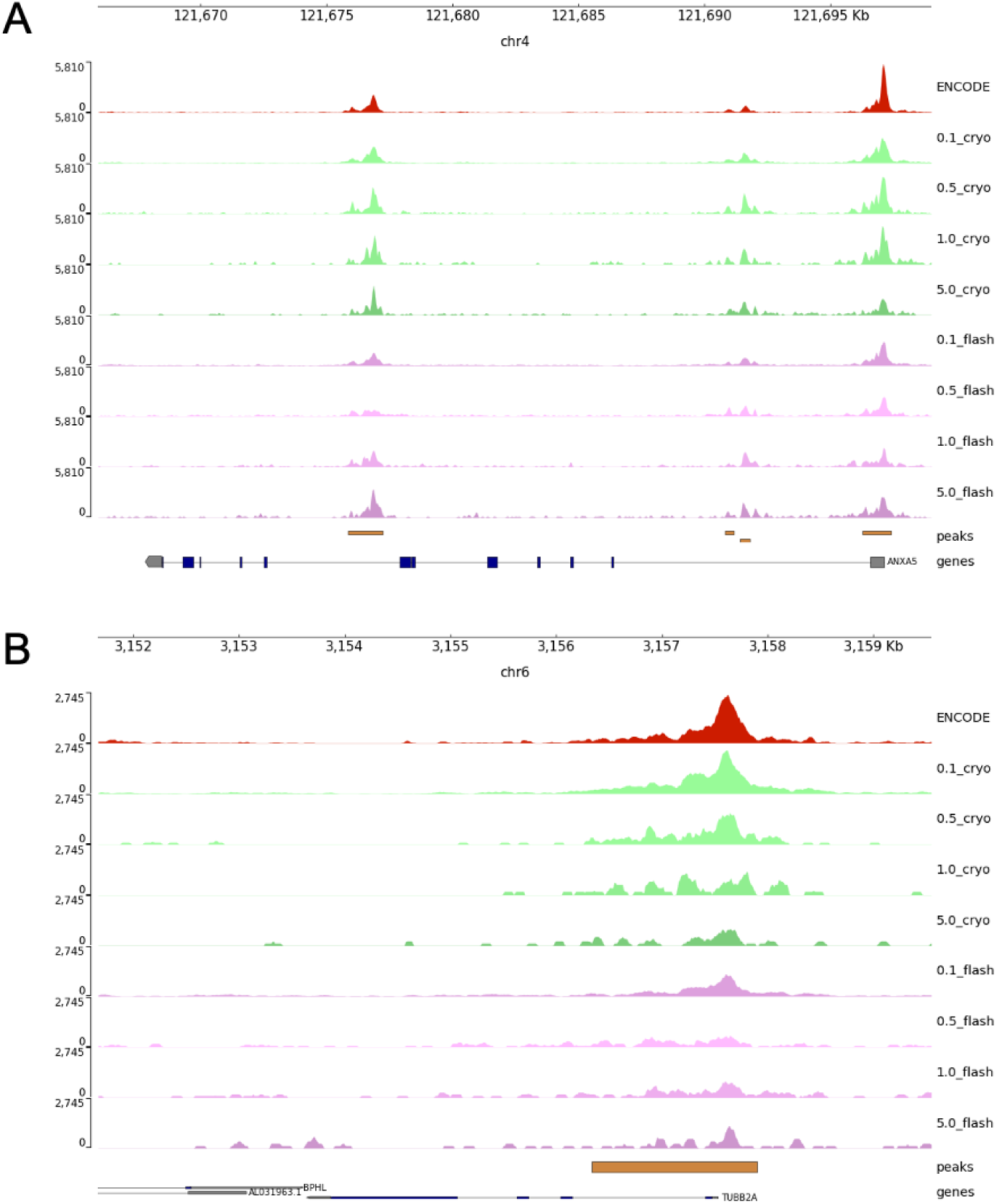
Genome tracks of selected regions. **a** Genome track of the ANXA5 gene. All protocols show similar signals around peak regions. **b** Genome track of the TUBB2A gene. While all protocols show signals within the peak region, the signal gets noisier with increasing FA concentration.

**Suppl. Fig. 2:**
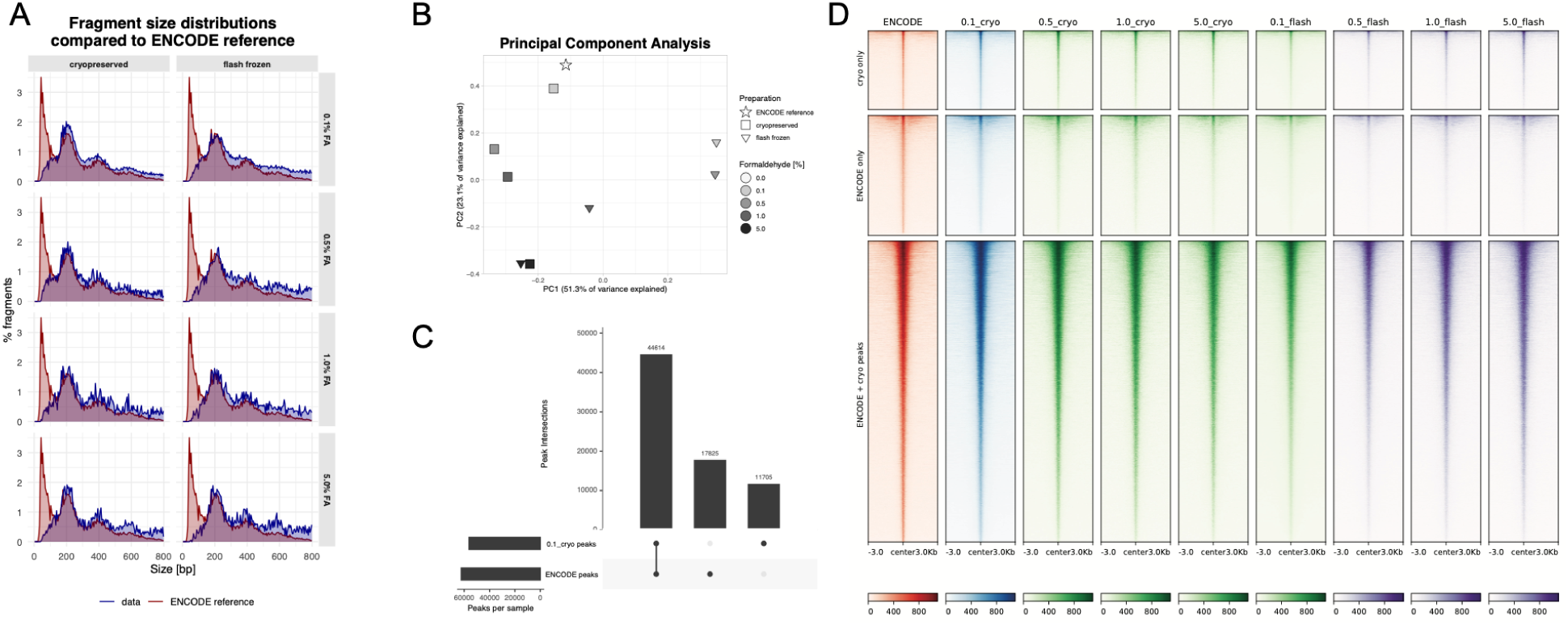
Further quality metrics on bulk ATAC-seq with sample preservation. **a** Fragment size distribution for all samples compared to a reference dataset. **b** PCA plot of all samples. **c** Upset plot showing the overlap between the reference peak set and a peak set called using the data from the best performing condition. **d** Heatmap showing peak sets found in c.

**Suppl. Fig. 3:**
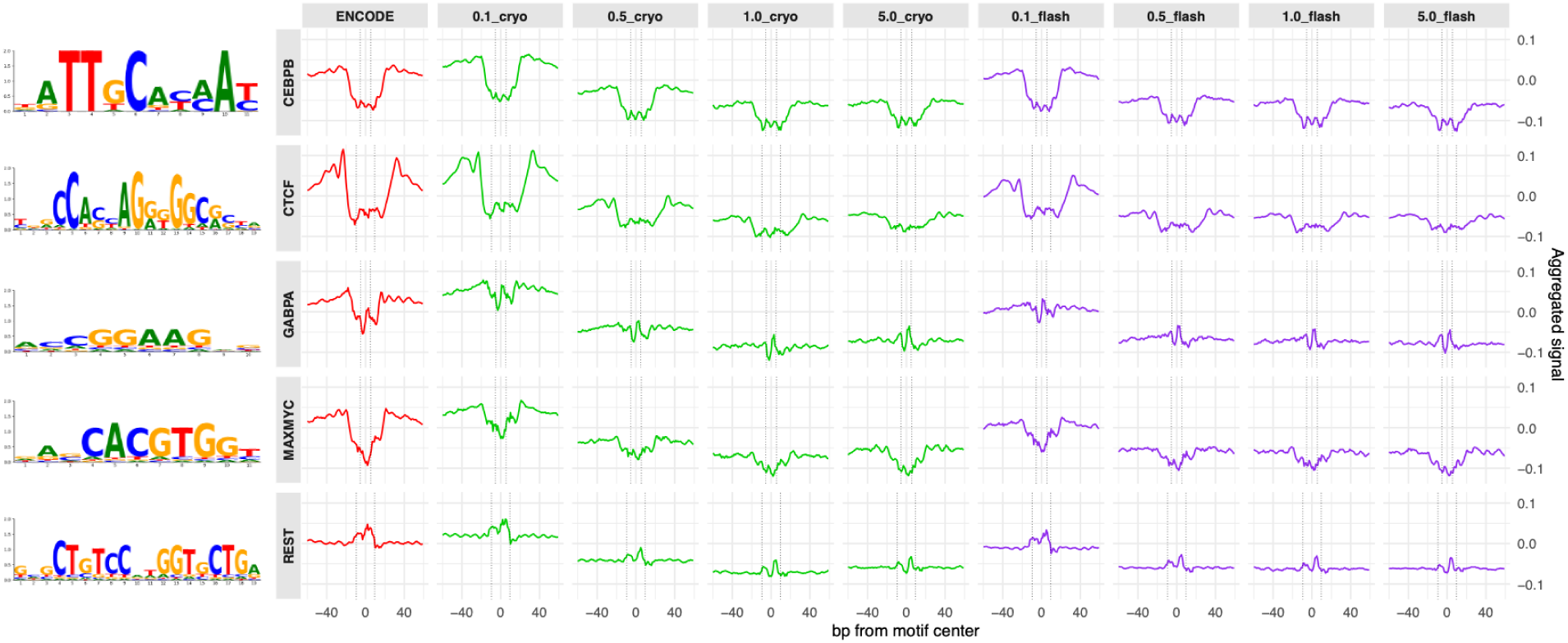
Footprint plots of selected transcription factors. Columns are samples, rows are transcription factors. Dotted lines show motif widths. While most footprints can be recovered for each sample, the strength decreases with increasing formaldehyde concentration. The GABPA footprint is visible only within the ENCODE and cryo_0.1 samples.

**Suppl. Fig. 4:**
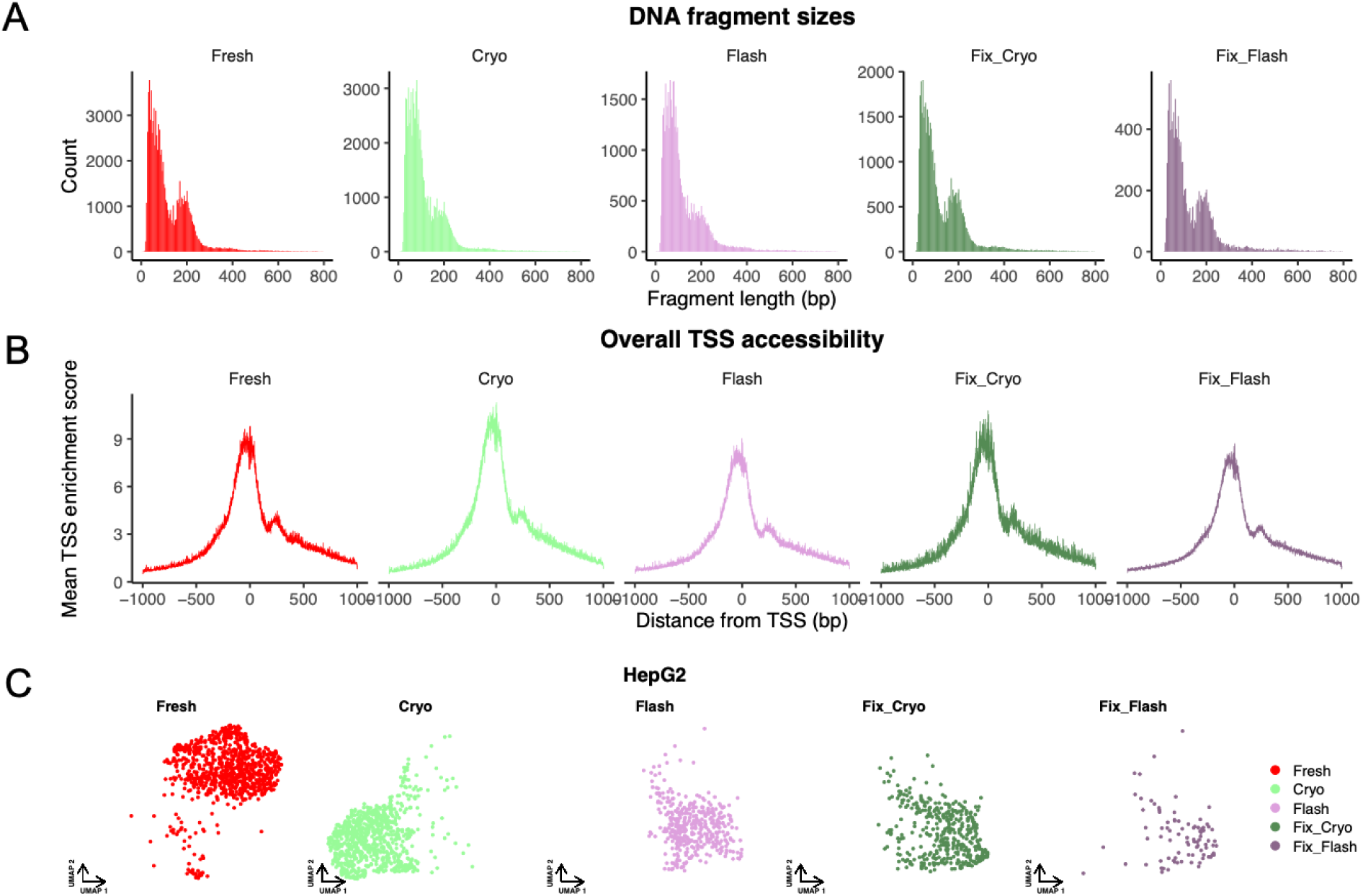
Quality control of HepG2 scATAC-seq data. **a** Per sample Fragment size histograms. **b** Mean TSS enrichment score around TSS per sample. **c** UMAP embedding split by sample.

**Suppl. Fig. 5:**
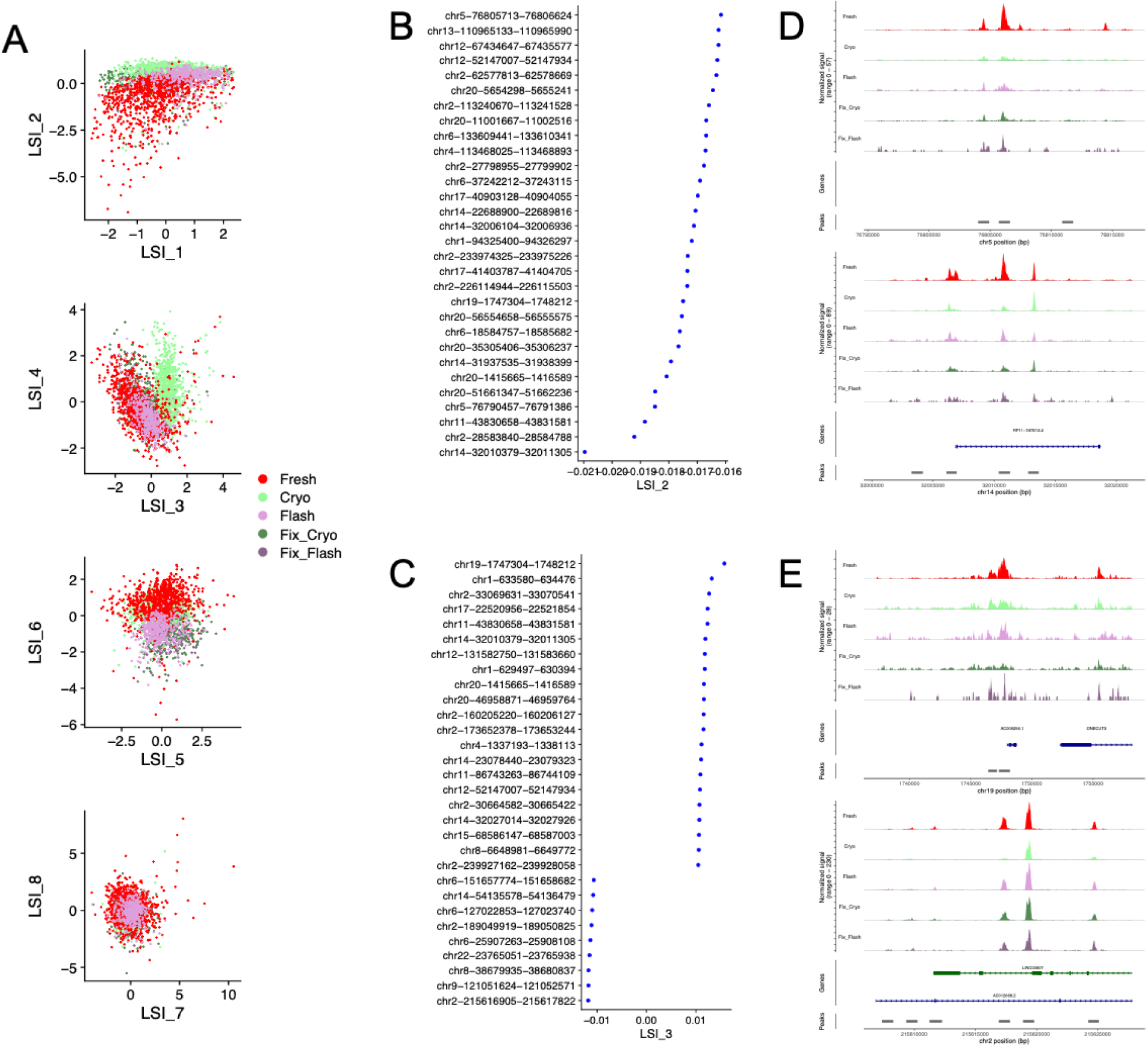
Investigating the LSI-based separation of samples on the UMAP embedding. **a** Scatterplots of several LSI components colored by sample. LSI 2 and 3 show the biggest sample separation. **b,c** Loadings of LSI components 2 and 3 visualizing the top associated regions. **d,e** Genome tracks top associated regions of LSI components 2 and 3.

**Suppl. Fig. 6:**
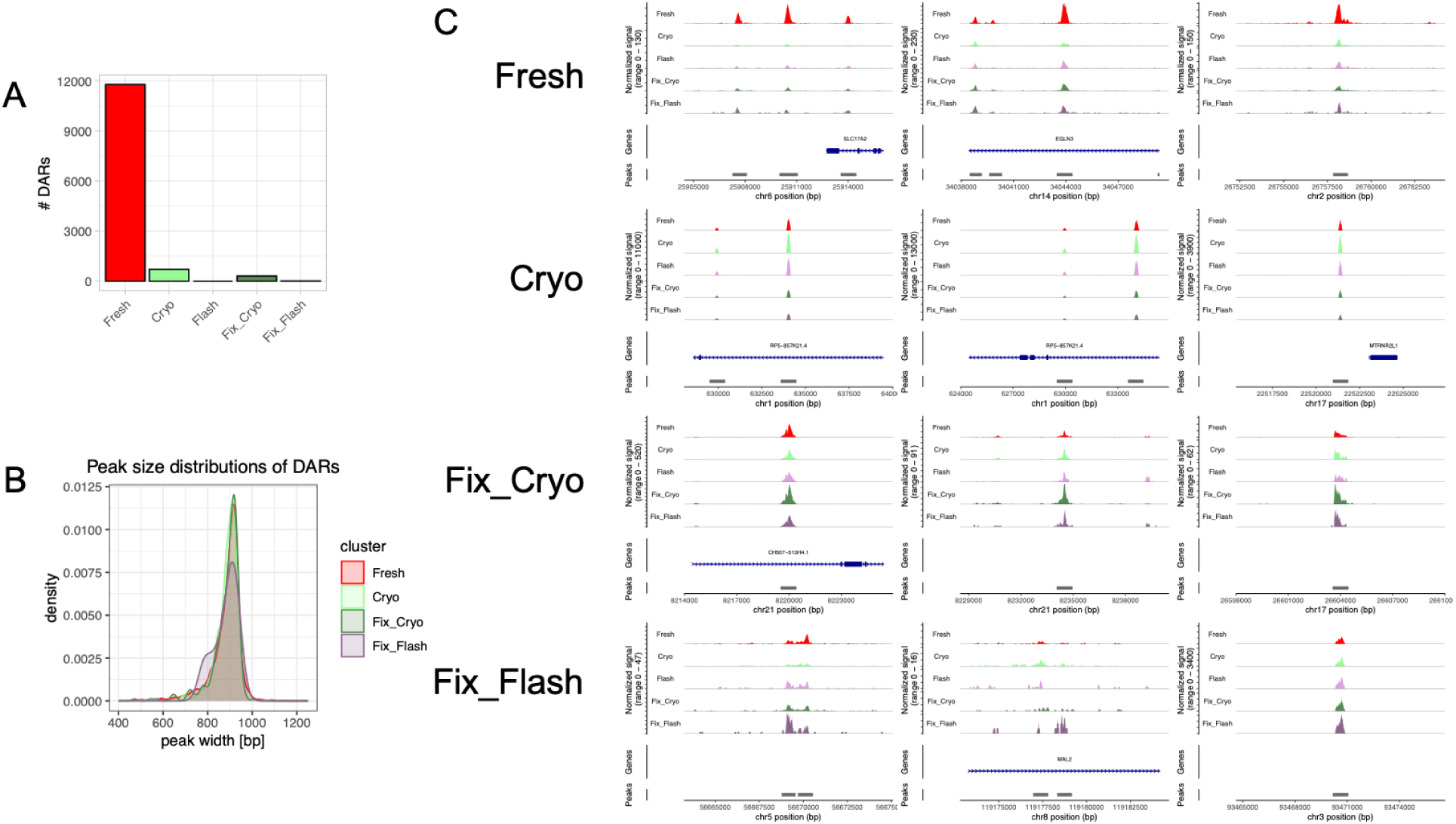
Differentially accessible peaks between conditions in scATAC-seq data. **a** Amount of DARs per sample. **b** Peak size distributions of DARs. **c** Genome tracks of top 3 DARs per sample. The Flash sample did not yield any DARs.

**Suppl. Fig. 7:**
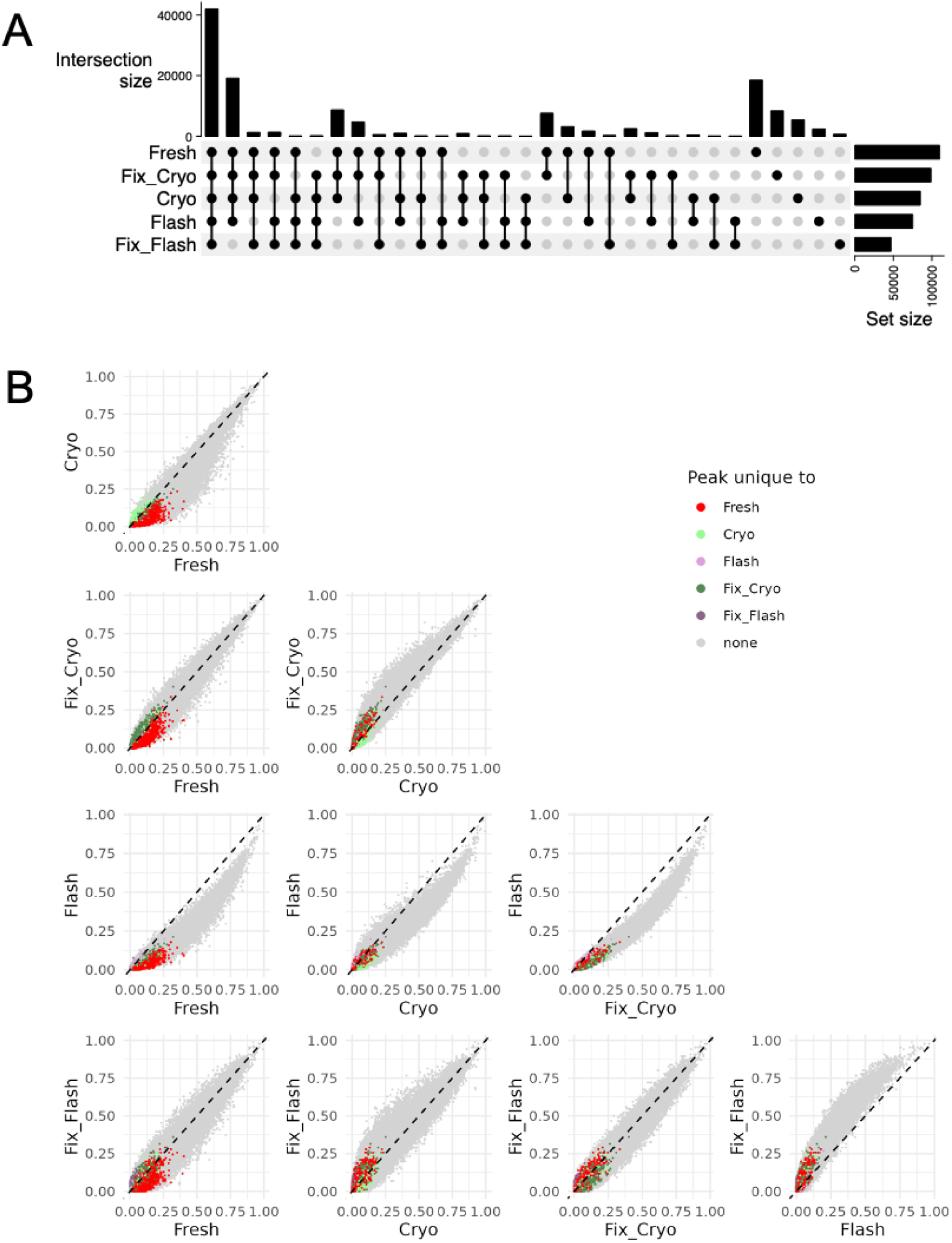
Sample specific peak calling. **a** Upset plot of called peaks over all samples. Only a small percentage of peaks are unique to a certain sample. **b** Correlation plots of peak appearance between samples. Each dot represents a peak of all peaks called per sample. The axes represent the fractions of cells within a certain sample that show at least one read within the respective peak region. Peaks are colored if they were only called in one sample.

**Suppl. Fig. 8:**
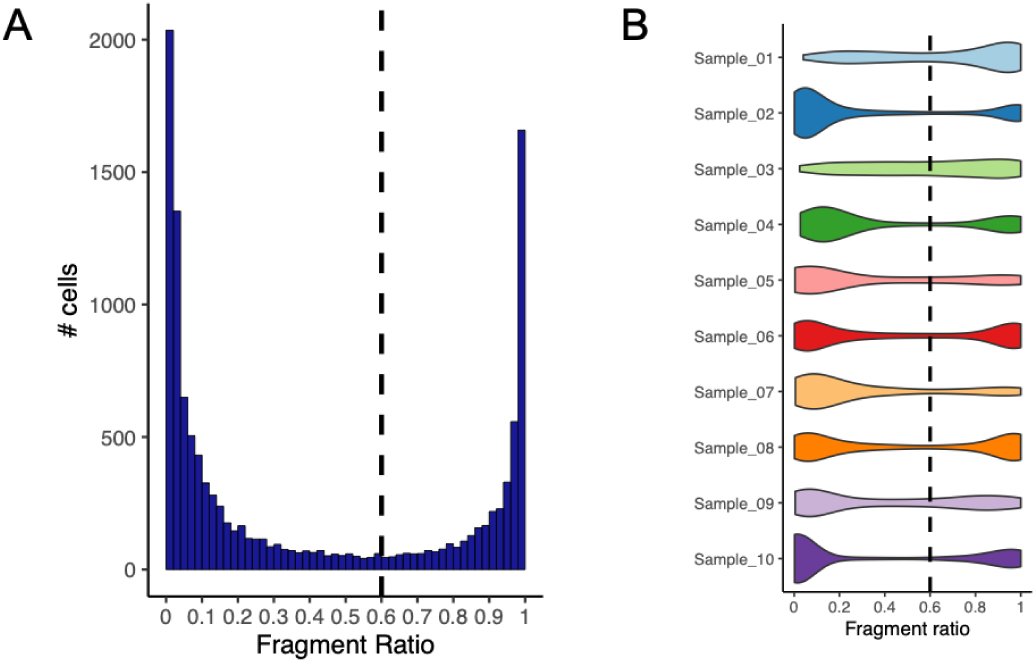
Sample barcode hopping in the context of MUX-scATAC-seq. **a** Histogram of fragment ratios for all sample and cell barcode combinations. **b** Violin plot of fragment ratios per sample for all sample and cell barcode combinations.

